# Will they survive? The critical status of Asiatic cheetahs in Iran

**DOI:** 10.64898/2026.11.16.766923

**Authors:** Atie Taktehrani, Mahsa Shah Hosseini, Navid Gholikhani, Kaveh Hobeali, Mohammad Hossein Karimi, Negin Samadzadeh, Hamed Abolghasemi, Ali Ranjbaran, Ahmad Radman, Ahmad Safarzadeh, Morteza Pourmirzai, Mohammad S. Farhadinia

## Abstract

The Asiatic cheetah (*Acinonyx jubatus venaticus*), once widespread, now survives only in Iran, with its population nearing extinction. In this study, we synthesised 13 years (2012–2024) of monitoring data to assess demographic and spatial patterns of this critically endangered subspecies in Iran. Using extensive camera trap surveys across eight reserves, supplemented by published records of cheetah occurrences on social media, we identified 24 adult cheetahs in the Northern and Southern Landscapes. No reproduction or new individuals have been recorded in the Southern Landscape for over a decade, while the Northern Landscape supports the remaining population, likely fewer than 30 adult individuals. From 2012 to 2024, at least 38 cubs were born from eight females in the Northern Landscape, with 74.1% surviving their first year. The absence of any newly detected adults throughout the 13-year study period suggests that long-term individual-based monitoring has likely captured the entire known adult population. Asiatic cheetahs show high mobility, often crossing unprotected communal lands and a major highway, increasing mortality risks. Camera traps enabled us to effectively identify individuals but are limited for tracking fine-scale movements, underscoring the need for satellite telemetry. Roadside fencing, wildlife underpasses, habitat protection, community-based land management, and stronger anti-trafficking measures are critical to enhance survival. Genetic risks, including low population size and inbreeding, further threaten viability. These findings inform conservation strategies for endangered carnivores in arid ecosystems.

## Introduction

Conserving small populations of large carnivores presents considerable challenges due to compounded genetic, demographic, and ecological constraints. In small and isolated populations, inbreeding and genetic drift can reduce individual fitness, lower juvenile survival rates, and increase disease susceptibility (Liberg et al., 2005; Hedrick & Fredrickson, 2010). Such genetic erosion is often driven by founder effects and population bottlenecks, which limit adaptive potential and heighten extinction risk, particularly when numbers fall below minimum viable population thresholds (Benson et al., 2019). These risks are especially pronounced in carnivores such as cheetahs (*Acinonyx jubatus*), which naturally exist at low densities and require extensive, connected habitats for viable population function, factors that complicate reliable population estimation (Becker et al., 2013; Belbachir et al., 2015).

In arid environments, cheetahs have to cope with low prey densities by adopting nomadic movement patterns to exploit patchy resources; however, this increases their exposure to human-wildlife conflict, including persecution and road mortality (Farhadinia et al., 2016a). Their large home ranges, often exceeding 500 km² in arid environments (Marker et al., 2008), combined with competition from larger predators (Pilaud et al., 2026), further threaten their survival. Male cheetahs, in particular, exhibit two distinct spatial tactics across their life history tied to reproductive strategy (Melzheimer et al., 2018; 2020). Males begin as floaters with extensive home ranges (averaging 1,595 km²) that overlap with those of females (mean 650 km²), before potentially establishing smaller resident territories (mean 379 km²) (Melzheimer et al., 2020).

Reproductive success in cheetahs is strongly limited by prey availability and habitat quality, with prolonged inter-birth intervals under resource scarcity constraining population growth (Wachter et al., 2011). Despite these ecological insights, population density and demographic structure remain poorly understood for many cheetah populations, particularly those inhabiting arid environments (Belbachir et al., 2015; Shams et al., 2025).

The Asiatic cheetah (*A. j. venaticus*), a critically endangered subspecies formerly distributed widely across West and Central Asia, has undergone a severe decline since the 1970s as a result of habitat loss, prey depletion, and increasing human–wildlife conflict (Farhadinia et al., 2017). Iran is now the only country with a known surviving population, making the Asiatic cheetah the rarest cheetah subspecies (Durant et al., 2024). Since 2001, published estimates have generally placed the known population at fewer than 70 individuals (Farhadinia et al., 2017; Khalatbari et al., 2023; Durant et al., 2024), distributed across several arid reserves in the eastern half of Iran (Ahmadi et al., 2017; Pilaud et al., 2026).

Despite more than two decades of conservation efforts in Iran (Farhadinia et al., 2017; Durant et al., 2024), the known Asiatic cheetah population remains very small and vulnerable and has not yet shown clear signs of recovery. Many historically occupied habitats have not yielded any confirmed cheetah detections in the past two decades, raising serious concerns about the subspecies’ long-term viability (Farhadinia et al., 2017; Khalatbari et al., 2023). This highlights the urgent need to understand the key demographic and spatial patterns shaping the survival of Asiatic cheetahs to guide current and future conservation strategies.

Since 2001, conservation efforts for the Asiatic cheetah have included decadal updates on its demographic and spatial patterns, primarily using photographic evidence (Farhadinia et al., 2016a; 2016b; 2017) or genetic markers (Khalatbari et al., 2023). In this study, we synthesised the most comprehensive dataset to date, spanning 13 years (2012–2024), to address: (1) What is the current state of Asiatic cheetahs in Iran? (2) What are the spatial patterns of Asiatic cheetahs? This synthesis provides critical insights into population persistence, distribution shifts, and emerging challenges, informing conservation priorities for this critically endangered subspecies.

## Study area

In Iran, cheetahs are primarily distributed across low-canopy rangelands characterized by sparse vegetation (density <25%) and rugged terrain, reflecting a habitat preference shaped by topographic complexity (Ahmadi et al., 2017). These environmental conditions share similarities with those inhabited by Saharan cheetahs in North Africa (Belbachir et al., 2015). Based on inter-reserve movement data (Farhadinia et al., 2016a) and the spatial distribution of reserves, the Asiatic cheetah population is divided into two subpopulations located around the central deserts of Iran, the Northern and Southern landscapes. A third subpopulation, Kavir, has yielded no confirmed cheetah detections in recent years (Farhadinia et al., 2017), raising concern that it may be locally extinct; however, its current status remains uncertain. This study focuses on the Southern and Northern landscapes, particularly the latter, which currently represent the most critical habitats for the subspecies.

The Northern Landscape, the primary breeding stronghold for Asiatic cheetahs in Iran, includes two surveyed reserves. The Southern Landscape encompasses 11 protected areas with confirmed cheetah presence, of which six were surveyed in this study (Table 1 and Fig. 1).

**Fig. 1.**
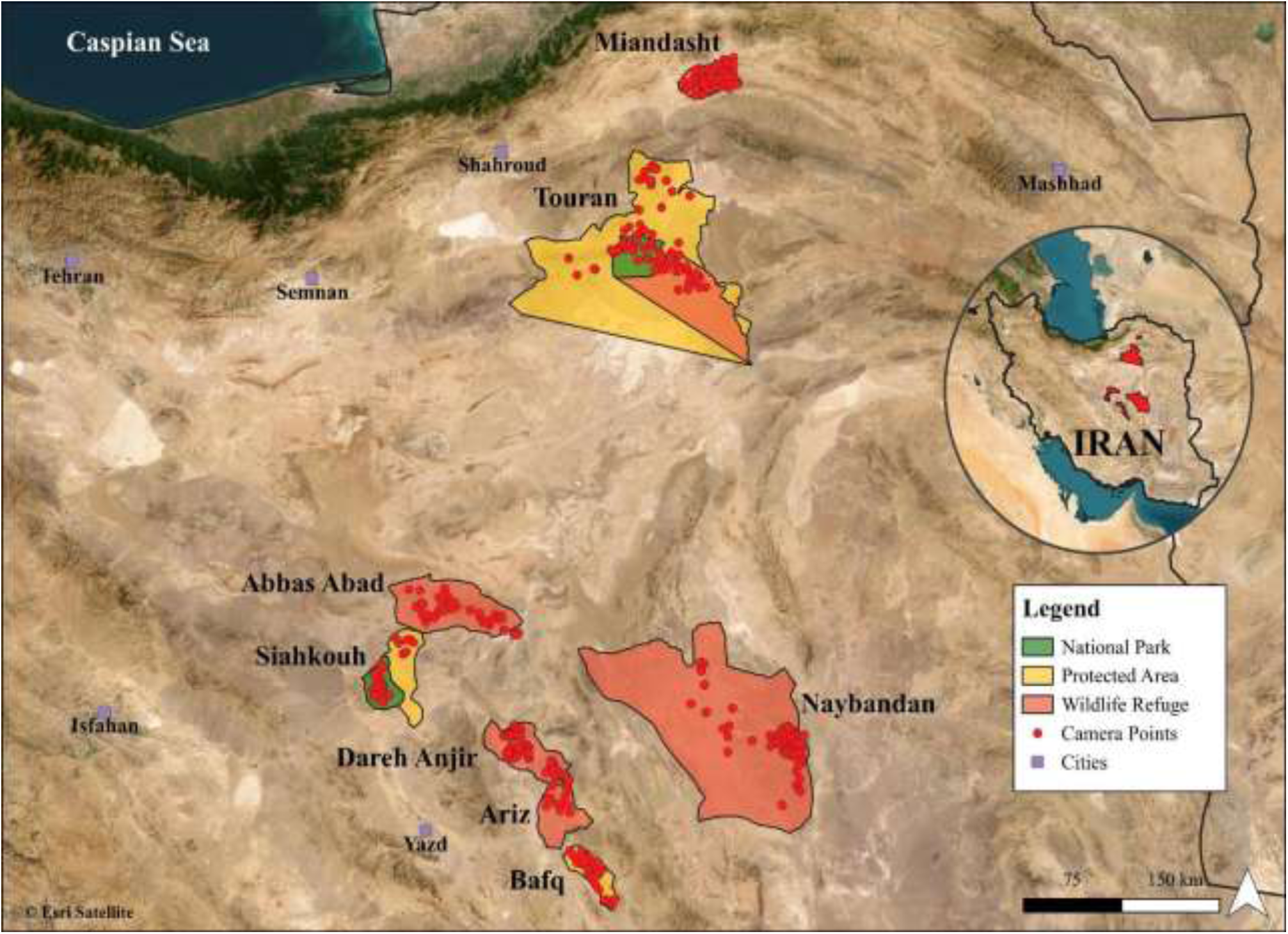
Map of the study area showing the spatial configuration of camera trap locations used for monitoring Asiatic cheetah (*Acinonyx jubatus venaticus*) populations in Iran. The map includes the Northern Landscape (Touran Biosphere Reserve and Miandasht Wildlife Refuge) and the Southern Landscape (Naybandan, Dareh Anjir, Ariz, and Abbas Abad Wildlife Refuges, as well as Siahkouh National Park, Siahkouh and Bafq Protected Area). Boundaries delineate the protected areas that were systematically monitored between 2012 and 2024

**Table 1.** Summary of systematic camera-trap surveys conducted across eight reserves in Iran during 2012–2023, including the number of cheetahs (with sex and cub status), the number of camera traps, the number of trap nights, the reserve area, and the year. M=male, F=female, C=cub and U=Unknown sex. PA=Protected Area, WR=Wildlife Refuge, NP=National Park and BR=Biosphere Reserve.

| Reserve/ Sampling number | Number of detected cheetahs | Number of camera trap stations | Sampling effort (Trap nights) | Area (km²) | Year |
|---|---|---|---|---|---|
| Siahkouh PA |  |  |  |  |  |
| Sampling 1 | 1F | 23 | 1051 | 2,000 | 2012 |
| Sampling 2 | 0 | 24 | 1600 |  | 2016 |
| Dareh Anjir WR |  |  |  |  |  |
| Sampling 1 | 4M | 26 | 1756 |  | 2012 |
| Sampling 2 | 3M | 22 | 1099 | 1,753 | 2015 |
| Sampling 3 | 3M | 26 | 1460 |  | 2016 |
| Bafq PA |  |  |  |  |  |
| Sampling 1 | 4 (3M &1U) | 47 | 2823 | 885 | 2011-2012 |
| Sampling 2 | 0 | 66 | 3901 |  | 2016 |
| Abbas Abad WR |  |  |  |  |  |
| Sampling 1 | 0 | 49 | 1889 | 3,054 | 2012 |
| Ariz WR |  |  |  |  |  |
| Sampling 1 | 3 (2M,1F) | 11 | 792 | 1310 | 2012 |
| Naybandan WR |  |  |  |  |  |
| Sampling 1 | 2 (1M & 1U) | 48 | 2662 |  | 2012-2013 |
| Sampling 2 | 1M | 33 | 461 | 15,160 | 2014-2015 |
| Sampling 3 | 2M | 50 | 2485 |  | 2016 |
| Miandasht WR |  |  |  |  |  |
| Sampling 1 | 4 (1F & 3C) | 32 | 2429 |  | 2012 |
| Sampling 2 | 5 (1M & 1F & 3C) | 16 | 1561 | 840 | 2013 |
| Sampling 3 | 3 (2M & 1F) | 30 | 1903 |  | 2014 |
| Sampling 4 | 7 (1M & 2F & 4C) | 8 | 272 |  | 2015 |
| Sampling 5 | 1M | 31 | 2135 |  | 2016 |
| Sampling 6 | 1M | 33 | 2067 |  | 2018 |
| Sampling 7 | 0 | 23 | 1465 |  | 2019 |
| Sampling 8 | 0 | 23 | 1611 |  | 2020 |
| Sampling 9 | 0 | 15 | 1814 |  | 2021 |
| Sampling 10 | 0 | 17 | 1476 |  | 2022 |
| Sampling 11 | 0 | 14 | 1775 |  | 2023 |
| Touran BR |  |  |  |  |  |
| Sampling 1 | 5 (3M & 2F) | 35 | 1176 |  | 2020 |
| Sampling 2 | 9 (2M & 3F & 4C) | 76 | 10177 |  | 2021 |
| Sampling 3 | 16 (2M & 6F & 8C) | 44 | 7702 | 14,000 | 2022 |
| Sampling 4 | 10 (3M & 3F & 4C) | 43 | 9574 |  | 2023 |

## Methods

### Camera trap surveys

We conducted systematic camera trap surveys across eight reserves in Iran between 2012 and 2024 (Table 1 & Fig. 4), supplemented by publicly available photographic records of Asiatic cheetahs shared by biologists, rangers, the Department of Environment, and citizen scientists on public platforms. Our surveys involved 865 camera trap stations, with a total sampling effort of 69,116 trap-nights (Fig. 1).

Due to the low density of cheetahs, optimal camera placement was challenging. Camera placement at scent-marking sites can produce male-biased detections (Brassine & Parker, 2015). To reduce this potential bias, we distributed cameras across a range of habitat features, including mountain passes, floodplains, open plains, and, where appropriate, water sources. Camera locations were selected using ranger knowledge and field observations. Water sources were treated as one of several potential locations of cheetah activity during warmer months(Pilaud et al., 2026), rather than as areas assumed to be used disproportionately by females with cubs (Laurenson, 1994; Farhadinia et al., 2016b).

We placed two cameras at crossing points to capture both flanks. We identified individuals by unique spot patterns; we excluded unidentifiable cheetahs. We determined sex visually by the presence of testicles in males.

### Reproductive and mortality assessment

We developed a photographic database to track cheetah families. Litter size was estimated from cubs first observed at 3–6 months of age. We classified a litter as lost when the female was subsequently detected without her cubs before they reached 17 months of age, the approximate period of maternal care until independence (Laurenson, 1994). Estimates of minimum observed survival to one year were restricted to litters with sufficient follow-up records to determine how many of the initially documented cubs were observed at or beyond one year of age. Mortality was considered confirmed only when supported by photographic evidence.

### Data analysis

We compiled cheetah detections into a presence–absence matrix across survey years. Unless death was confirmed, we assumed individuals were alive until their last photographic record. Duration of detection (years) was calculated for each cheetah, with median and interquartile range (IQR) reported to minimize outlier influence, given the small sample size.

Our spatial analyses focused on the Northern Landscape during 2020–2024, when continuous, systematic camera-trap monitoring provided sufficiently consistent location records for identified individuals to allow a more reliable characterization of their observed ranging patterns. Camera-trap detections were supplemented by a small number of documented records from rangers. Available records from the Southern Landscape during this period were insufficient for spatial analysis, and the ranging patterns of individuals previously recorded there had already been characterized in earlier research (Farhadinia et al., 2016a). We processed detection points in QGIS 3.40.4 to identify unique individuals and estimated movement ranges using the Minimum Convex Polygon (MCP) method (Börger et al., 2006). Only individuals with ≥7 detections were included to ensure reliable estimates.

We used annual counts of cheetahs detected from photographic evidence collected between 2014 and 2024 across the Northern Landscape. Although monitoring began in 2012, the GAM analyses were restricted to 2014–2024 because the earlier records were less consistent in coverage and data quality. To describe temporal patterns in detected demographic metrics, we fitted generalized additive models (GAMs) in R using the mgcv package. GAMs accommodate non-linear relationships between predictors and response variables, with effective degrees of freedom (edf) indicating the complexity of the fitted smooth (Wood, 2017).

For each metric, including adult males, adult females, cubs of the year (<1 year), total adults, total detected individuals, and mortality, we fitted a GAM with a smooth effect of year and a negative binomial error distribution to account for overdispersed count data. Model predictions with 95% confidence intervals were generated for each year. Model fitness and assumptions were carefully evaluated to ensure adequate flexibility without overfitting. Because the models did not explicitly account for variation in sampling effort or detection probability, the results were interpreted as exploratory trends in detected individuals rather than estimates of changes in true population size.

## Results

### Demographic patterns

Over the 13-year study period, we identified a total of 24 adult Asiatic cheetahs in Iran (Fig. 2), comprising 14 males, 9 females, and one individual of unknown sex. We recorded 15 individuals in multiple reserves and unprotected areas, indicating high inter-reserve movements. The highest number of individuals was detected in Touran Biosphere Reserve (11 cheetahs: 6 females, 5 males). Miandasht Wildlife Refuge accounted for five individuals, while eight cheetahs were recorded across the Southern Landscape (Dareh Anjir Wildlife Refuge, Siahkouh National Park, Siahkouh Protected Area and Naybandan Wildlife Refuge). Notably, observed sex ratios varied across landscapes, from male-biased in the Southern Landscape (6:1) to balanced in the Northern Landscape (8:8) (Fig. 2), which may reflect underlying spatial differences in population structure and reproductive dynamics, though random variation across such small samples cannot be excluded.

**Fig. 2.**
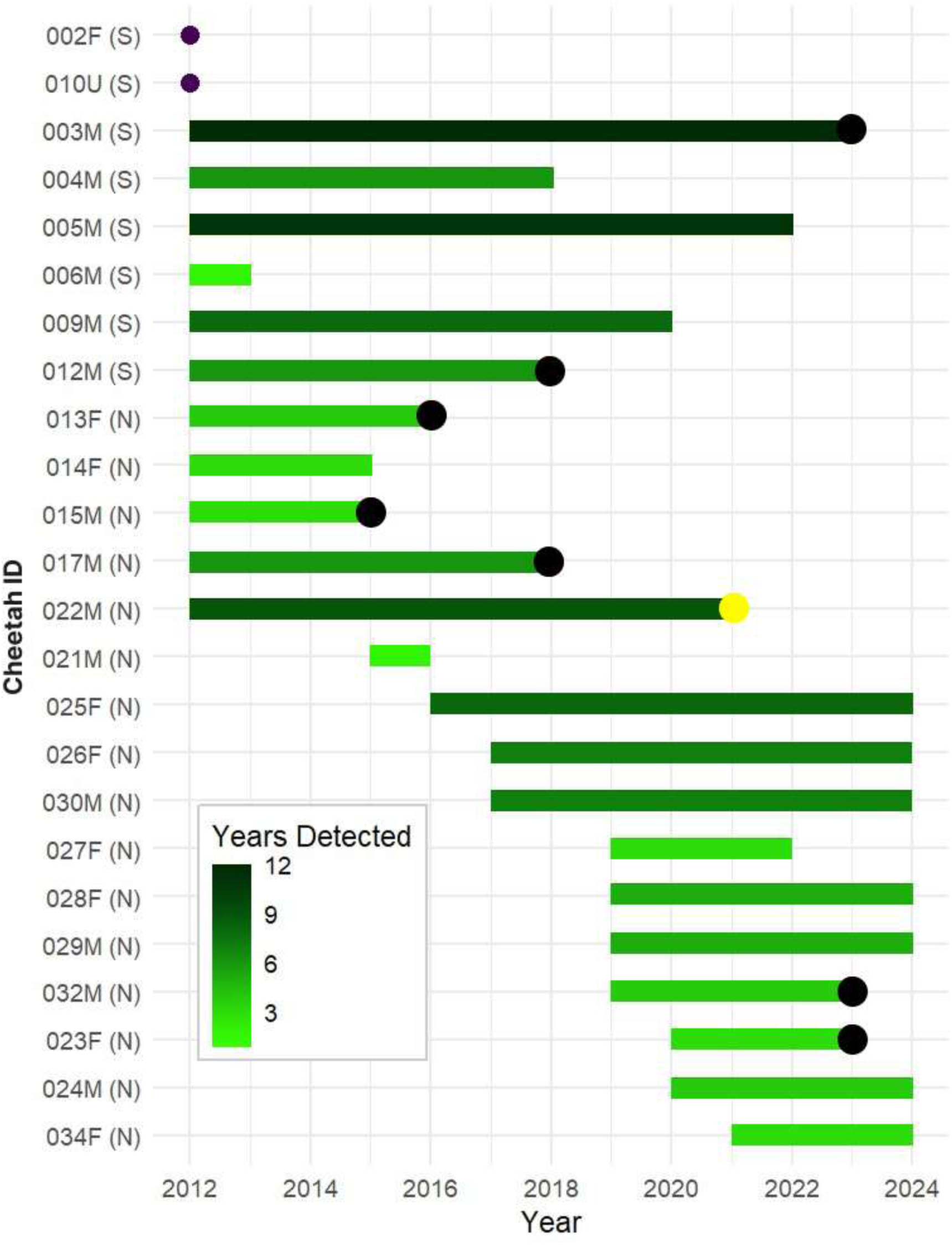
Abacus plot illustrating the annual detections of individual Asiatic cheetahs in Iran (2012– 2024). Each bar represents one cheetah (N = Northern Landscape; S = Southern Landscape), with length indicating years detected. Black circles denote confirmed mortality; yellow circles indicate capture. This highlights temporal patterns, population persistence, and high turnover in the critically endangered subspecies.

The median duration of camera-trap detections for individual cheetahs was 5.5 years (IQR = 4), with males showing a longer presence (Median = 7, IQR = 5) compared to females (Median = 4, IQR = 3). Since all individuals were recorded as adults, some likely reached the end of their natural lifespan during the study. Between 2020 and 2024, systematic monitoring in the Northern Landscape identified 11 cheetahs, of which 3 either died or were removed from the wild, leaving at least 8 adults in the population by the end of 2024. In the Southern Landscape, no new cheetah individuals have been recorded since 2011. The last known cheetah in the Southern Landscape was found dead in June 2023, and there have been no further detections since then (Fig. 2).

The last cheetahs known to be born in the Southern Landscape were a coalition of three brothers (003M, 004M, 005M) in 2010, with the last known female recorded there in 2012. In contrast, all documented breeding events since 2010 have occurred exclusively in the Northern Landscape, where known individuals have been repeatedly detected. In recent years, no previously unrecorded adults have been identified that would suggest breeding outside the documented family groups (Fig. 2).

Between 2012 and 2024, we identified 14 cheetah families in the Northern Landscape, with a mean litter size of 2.7 ± 1.1 cubs (SD) when first detected at 3–6 months of age. These families represented litters produced by eight females, with 38 cubs documented in total. All 10 families that were subsequently recorded had at least one cub observed accompanying its mother at or beyond one year of age. Across these families, the proportion of initially documented cubs observed accompanying their mothers at or beyond this age corresponded to a minimum observed survival of 74.1%. Within these families, cubs not subsequently observed with their mothers at or beyond one year of age were either confirmed dead or no longer detected, and their fate could not always be determined. Four of eight females produced multiple litters during the monitoring period (Table 2). Because some cubs may have survived but remained undetected, and families that were not subsequently recorded were excluded from the calculation, this proportion should be interpreted cautiously and should not be generalised to all cubs in the population.

**Table 2.** Cubs born, last recorded sightings, and known cub losses for Asiatic cheetah females in Miandasht Wildlife Refuge and Touran Biosphere Reserve, Iran, during 2012–2024. In the column “Cubs at 3–6 months (cubs at 1 year),” the first value indicates the number of cubs detected at 3–6 months of age, and the value in parentheses indicates the number subsequently detected at or beyond one year of age. An asterisk (*) identifies the 10 litters monitored through the cubs’ first year of life and included in the estimation of minimum observed cub survival to one year. The cubs of females 026F and 034F were recorded at one year of age in 2025; these follow-up records are included to document the outcome of a litter first recorded during the study period. NA (not available) indicates that the information specified in the corresponding column was not recorded.

| **Reserve** | **Individual<br>ID** | **cubs 3-6m<br>(cubs 1y)** | **Birth year** | **Last seen<br>with mother** | **Cub presumed<br>lost (<1 year)** | **Remarks** |
|---|---|---|---|---|---|---|
| Miandasht WR | 013F | 3 (3)<sup>*</sup> | 2012 | Sep 2013 | - | - |
|  |  | 1 (1)<sup>*</sup> | 2015 | Nov 2015 | - | - |
|  | 014F | 3 (NA) | 2015 | Oct 2015 | NA | - |
| Touran BR | 025F | 3 (2)<sup>*</sup> | 2020 | May 2020 | NA | - |
|  |  | 2 (1)<sup>*</sup> | 2021 | Feb 2022 | 1Cub lost | Mother subsequently recorded without the cub |
|  |  | 2 (2)<sup>*</sup> | 2023 | Aug 2024 | NA | - |
|  | 026F | 3 (2)<sup>*</sup> | 2020 | Dec 2020 | 1 Cub lost | Mother subsequently recorded without the cub |
|  |  | 2 (NA) | 2021 | Aug 2021 | NA | - |
|  |  | 4 (2)<sup>*</sup> | 2023 | Jan 2025 | 2 Cubs lost | Mother subsequently recorded without cubs |
|  | 027F | 4 (NA) | 2022 | Sep 2022 | NA | - |
|  | 028F | 4 (2)<sup>*</sup> | 2022 | June 2023 | 2 Cubs lost | Mother subsequently recorded without 2 cubs |
|  |  | 2 (NA) | 2024 | NA | NA | - |
|  | 023F | 1 (1)<sup>*</sup> | 2021 | Aug 2022 | - | - |
|  | 034F | 4 (4)<sup>*</sup> | 2024 | 2025 | NA | - |

Between 2012 and 2024, 21 losses from the wild cheetah population were recorded, comprising deaths and removals from the wild, with all but two occurring in the Northern Landscape (Table S1). The leading cause was road collisions (8 deaths), followed by human-wildlife conflict (4 deaths). Other causes included orphaned cub removals (3), unknown factors (3), and single cases of illegal trade of cubs, veterinary malpractice, and natural causes. Confirmed cases of cubs being removed from the wild by humans (n=4) include one individual trafficked for the illegal wildlife trade (2017), two females taken by a shepherd (2022), and one individual captured due to apparent malnourishment (2024; Table S1).

The GAM indicated a significant temporal trend in the number of detected cheetahs from 2014 to 2024 (edf = 1.47, χ² = 11.75, p = 0.006), explaining 70.5% of the deviance (adjusted R² = 0.58), with a moderately non-linear pattern over time (Fig. 3). This result indicates variation in the number of detected individuals over time but does not by itself demonstrate population growth. In contrast, the temporal trend in detected mortality was not significant (edf = 1.76, χ² = 1.50, p = 0.46). The mortality model explained 23.2% of the deviance (adjusted R² = 0.11), indicating limited explanatory power. The small number of annual observations may have limited the ability to detect temporal variation in mortality.

**Fig. 3.**
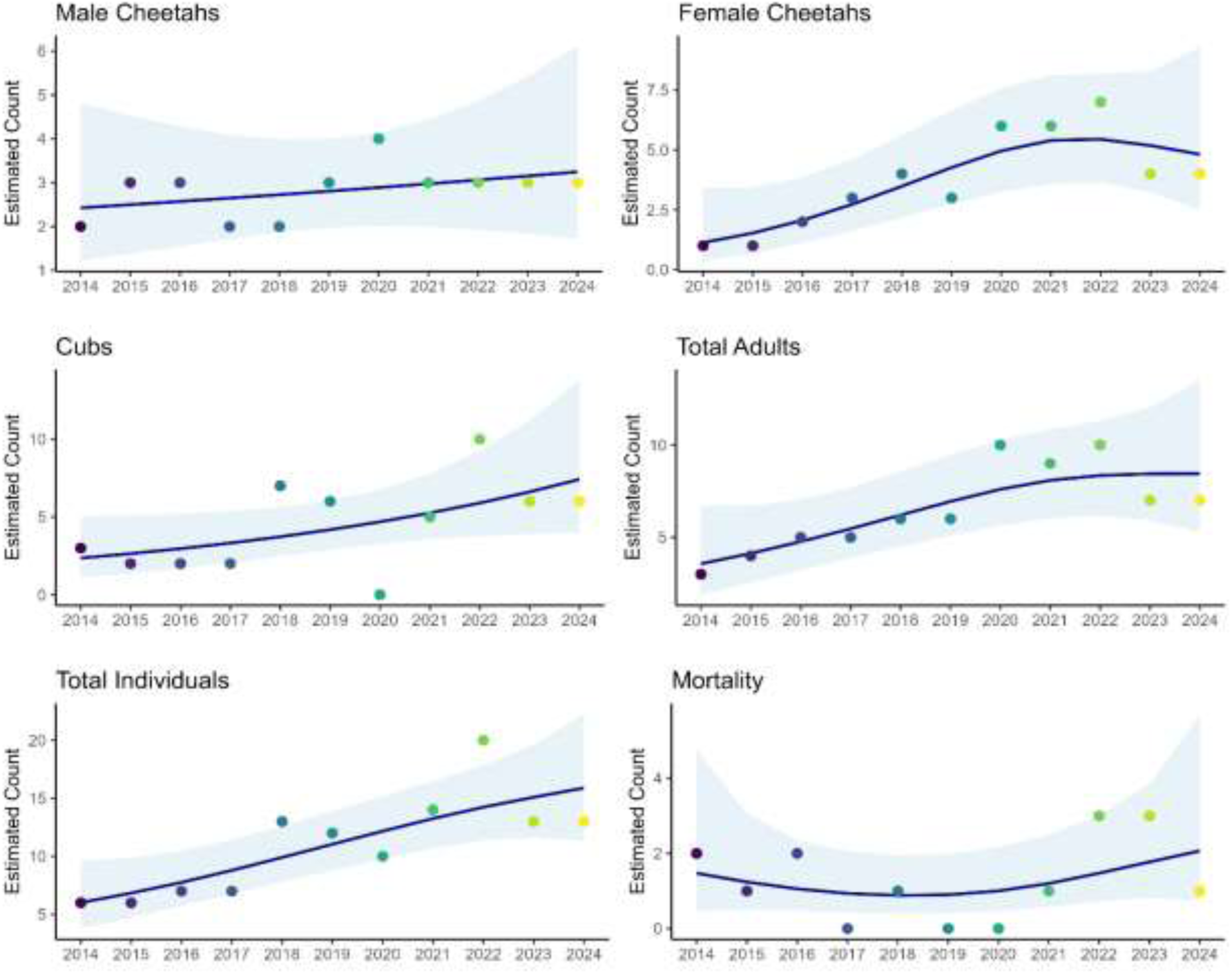
Temporal trends in Asiatic cheetah demographic metrics from 2014 to 2024 across the Northern Landscape in Iran. Each panel displays estimated counts for demographic parameters, including males, females, cubs, total adults, total individuals, and mortality, modelled using generalized additive models (GAMs) with a negative binomial error distribution. Solid lines show model-predicted trends, shaded ribbons represent 95% confidence intervals, and points indicate observed annual counts. Years are shown on the x-axis. Note that counts reflect detected individuals based on photographic evidence and may not represent the true population size.

**Fig. 4.**
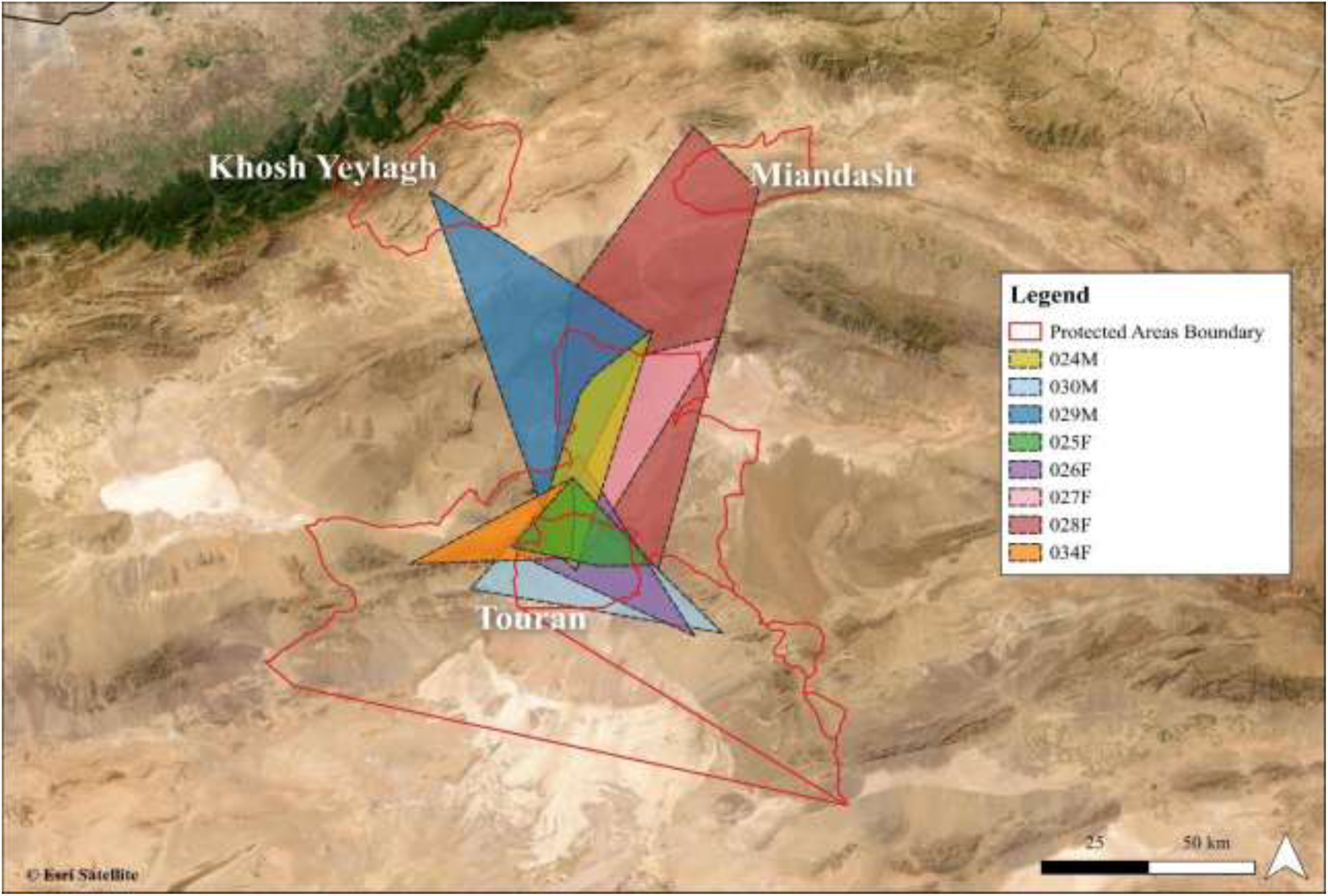
Recent ranging patterns of identified cheetahs in the Northern Landscape of Iran during 2020–2024, based primarily on camera-trap detections, supplemented by occasional documented records from rangers. Each coloured minimum convex polygon (MCP) represents the observed range of the corresponding cheetah identified in Figure 2. Detections recorded before 2020 were excluded to improve map clarity and focus on recent ranging patterns.

Although monitoring began in 2012, the GAM analyses were restricted to 2014–2024 because the earlier records were less consistent in coverage and data quality.

### Spatial patterns

The median maximum distance between the farthest recorded locations for Asiatic cheetahs in the Northern landscape was 81.9 km (IQR: 40.3 km). The median maximum distance was greater for males than for females (males: 83.1 km, IQR: 39.5 km; females: 70.6 km, IQR: 67.6 km). Similarly, the median MCP area for individual cheetahs was 1,354 km² (IQR: 1,514 km²), with larger ranges for males (1,524 km², IQR: 2,021 km²) than females (1,184 km², IQR: 1,354 km²). The spatial overlap of male and female cheetahs’ roaming areas, based on five years of camera trap data and documented rangers’ records (2020–2024), is illustrated in Fig. 4.

Of the eight identified Asiatic cheetahs, four individuals (025F, 026F, 030M, and 034F) were exclusively recorded within the Touran Biosphere Reserve. The remaining four (024M, 027F, 028F, and 029M) were initially detected in Touran but later observed in adjacent communal lands or other reserves (Table 3). Individual 027F, after separating from her mother and producing her first litter, was never seen again in Touran; all subsequent sightings occurred in unprotected communal lands. The location data for this female are from when she was within the Touran Biosphere Reserve (Fig. 4), thus providing limited information on her movement range outside the Touran.

**Table 3.** Spatial patterns of Asiatic cheetah movement across the Northern Landscape of Iran during 2020–2024.

| Cheetah ID | Max distance between farthest locations (km) | MCP (km²) | Reserves crossed |
|---|---|---|---|
| 024M | 83.1 | 1038 | Touran BR, communal land |
| 025F | 45.1 | 629 | Touran BR |
| 026F | 70.6 | 1184 | Touran BR |
| 027F | 89.7 | 1694 | Touran BR, communal lands |
| 028F | 149.3 | 5687 | Touran BR, communal land, Miandasht WR |
| 029M | 120.2 | 3059 | Touran BR, communal land, Khosh Yeilagh WR |
| 030M | 80.7 | 1524 | Touran BR |
| 034F | 58.6 | 686 | Touran BR |

Individual 028F displayed the widest movement range (Fig. 4): born in Touran, she gave birth to her first litter in communal lands, returned to Touran with her cubs, and later moved her second litter northward to Miandasht, ultimately covering 5,687 km², the largest MCP area recorded in this study. Similarly, individual 029M, also born in Touran, went undetected for three years before being re-sighted in both the Touran Biosphere Reserve and Khosh Yeilagh Wildlife Refuge.

## Discussion

Over the 13-year study period (2012–2024), only 24 adult Asiatic cheetahs were detected in Iran, despite extensive survey efforts totalling 69,116 trap nights across 27 distinct sampling sessions in eight reserves (Table 1). Seven of the 24 individually identified adult cheetahs (29.2%) were confirmed dead by the end of 2024. No evidence of Asiatic cheetah presence remains in the Southern Landscape as of 2024, with all recent detections confined to the Northern Landscape. Given the small sample size, we caution against drawing firm conclusions about population trends over the past 13 years. However, collectively, these findings indicate an extremely small population, raising serious concerns about the subspecies’ long-term survival in the wild.

### Demographic patterns

Our data show that the Northern Landscape remains the only documented breeding stronghold for the Asiatic cheetah. In years with recorded breeding, 1–3 females produced documented litters comprising a total of 3–8 cubs annually (Table 2). Between 2012 and 2024, 38 cubs were recorded in this landscape, and a substantial proportion (74.1%) were observed beyond one year of age, indicating continued reproduction and some potential for recruitment. However, this reproduction has not yet resulted in a clear increase in the number of detected adults. This may reflect slow recruitment into the adult population, mortality after the first year, dispersal, or incomplete detection in areas not covered by our monitoring efforts.

The observed cub survival in Iran falls within the broad range reported for African cheetah populations, although direct comparisons are limited by differences in monitoring effort, follow-up periods, and survival definitions. In the Northern Landscape, all ten monitored families had at least one cub survive to its first year, similar to the pattern observed during the 2010s, when at least one cub from every monitored family survived this milestone (Farhadinia et al., 2016b). The estimated cub survival beyond one year was 74.1%, which appears lower than the 89.5% reported for Iran during the 2010s (Farhadinia et al., 2016b), suggesting a possible decline in cub survival. However, this difference should be interpreted cautiously because differences in monitoring effort, data availability, and small sample sizes may exaggerate the apparent contrast between periods. For comparison, reported cub survival from 4 to 14 months ranges from 54.5% in the Serengeti to 95.8% in the Kgalagadi Transfrontier Park (Laurenson, 1994; Mills & Mills, 2014). The Northern Landscape estimate therefore falls within the range reported for African cheetahs, although differences in study design limit direct comparison.

Losses among Asiatic cheetah cubs varied among years, but their causes remain poorly understood. In African populations, cub mortality is often associated with predation and starvation (Laurenson, 1994; Mills & Mills, 2014) whereas anthropogenic causes contributed substantially to documented mortality in Iran. Records of losses from the wild population between 2012 and 2024 (Table S1) include 21 cheetahs that died or were removed from the wild, 19 in the Northern Landscape and two in the Southern Landscape. Because recent breeding has been concentrated in the Northern Landscape, losses there affected the remaining reproductive core, including females and cubs, with potential consequences for recruitment.

Earlier assessments (2001–2016) identified human–wildlife conflict as one of the main documented causes of cheetah mortality, followed by road collisions and poaching (Farhadinia et al., 2016b; 2017). In our more recent records, road collisions were the most frequent documented cause of mortality. However, recent trends suggest an apparent reduction in conflict-related mortality, likely due to awareness and education campaigns, while road-related deaths have increased, possibly driven by expanding infrastructure (Mohammadi et al., 2023).

Although the GAM indicated an increase in the number of detected cheetahs over time, this pattern should not be interpreted as evidence of population growth. The analysis did not explicitly account for variation in camera-trap coverage, survey effort, or detection probability. The apparent increase may therefore reflect improved detection resulting from expanded camera coverage, better knowledge of cheetah locations, or improved field methods. Accordingly, the GAM results are best interpreted as describing temporal variation in detected individuals rather than changes in the true population size. Given the small number of annual observations and the limitations of the available data, no firm conclusion can be drawn about the population trend during 2014–2024.

### Spatial patterns

Asiatic cheetahs demonstrated high spatial mobility across the drylands of the Northern Landscape, often traversing multiple reserves and unprotected communal lands. Of the eight adult cheetahs identified in the Touran Biosphere Reserve, four moved into neighbouring landscapes. The median maximum distance between the farthest recorded detection points for individuals in the Northern Landscape was 81.9 km (IQR: 40.3 km), with males generally travelling farther than females; the maximum recorded distance was 149.3 km for female individual 028F. In the Southern Landscape, multiple males were previously recorded at locations more than 150 km apart, with one male travelling up to 400 km between reserves (Farhadinia et al., 2016a). Similar long-distance movements have been observed in other arid regions: in Ahaggar Cultural Park, Algeria, the mean maximum distance travelled by two male cheetahs was approximately 45 km (Belbachir et al., 2015), while longer distances were also documented for males in the semi-arid farmlands of Namibia (Marker et al., 2008; Melzheimer et al., 2020). The high mobility of Asiatic cheetahs and their large spatial requirements may also serve as a mechanism for inbreeding avoidance (Khalatbari et al., 2023).

Similarly, the median MCP area for individual cheetahs in the Northern Landscape was 1,354 km², generally smaller than estimates from the 2010s, which were mostly recorded in the Southern Landscape (2,105.3 ± SE 778.6 km²;(Farhadinia et al., 2016a)). While both the current study and Farhadinia et al. (2016a) rely on camera trap data spread across expansive drylands, the estimated ranges remain considerably larger than those derived from the only GPS telemetry data available for Asiatic cheetahs (1,137 km²; (Cheraghi et al., 2018)). This highlights the species’ extensive spatial requirements and high mobility in arid environments. Comparable MCP estimates have been reported only in Namibia, where cheetahs in semi-arid farmlands occupy home ranges between 1,344 and 2,863 km² (Marker et al., 2008; Melzheimer et al., 2020). In contrast, cheetah ranges in sub-Saharan Africa are typically smaller (e.g., Houser et al. (2009)), although Belbachir et al.(2015) reported a camera trap-based MCP estimate of 1,337 km² for Saharan cheetahs, reinforcing the pattern of large home ranges in more arid ecosystems.

. All individuals included in the spatial analysis were either detected in or originated from the Touran Biosphere Reserve, highlighting its central importance as a stronghold for the remaining Asiatic cheetah population. However, individuals are frequently dispersed beyond reserve boundaries, often crossing paved highways that fragment the landscape. These roads, while not impeding movement, significantly increase mortality risk, particularly from vehicle collisions, which have become a leading cause of death (Mohammadi et al., 2018; 2023).

### Conclusion and management recommendations

Our analysis indicates that fewer than 30 adult Asiatic cheetahs are currently known in Iran. This number is lower than earlier broad estimates of approximately 50 individuals (Farhadinia et al., 2017), although differences in survey methods limit direct comparison. The current demographic status is also consistent with recent genetic assessments reporting an effective population size of 11–17 individuals (Khalatbari et al., 2023), although effective population size should not be interpreted as a direct count of wild cheetahs. At such critically low numbers, the population remains highly vulnerable to stochastic events, genetic deterioration, and demographic collapse.

Across all monitoring years, no more than 11 adult individuals were detected annually, alongside an estimated 3–8 dependent cubs per year. The absence of any newly detected adults throughout the 13-year study period suggests that long-term individual-based monitoring has likely captured the entire known adult population. However, incomplete spatial and temporal coverage across the Northern Landscape, due to its large extent and limited camera-trap effort, likely resulted in intermittent non-detection of known individuals and consequent underestimation of individual longevity.

The temporal analyses provide an exploratory description of changes in detected individuals, but they do not establish a biological increase in population size. Conservation recommendations in this study are therefore based primarily on observed demographic and spatial patterns, including documented losses from the wild population, rather than on evidence of population growth from the GAMs.

Drawing on both the findings of this study and the broader conservation literature, we propose the following conservation recommendations:

1. The critically small known population and evidence of low genetic diversity threaten long-term viability (Khalatbari et al., 2023). Although our study provides further evidence of the small population, it does not include new genetic analyses. Therefore, before considering genetic rescue through admixture with African cheetahs which was previously suggested (Khalatbari et al., 2023), a current genetic profile of the population should be developed, including pedigree relationships, to supplement previous genetic research. Given that the Iranian Department of Environment is implementing a captive-breeding programme, the removal of cubs or subadults from the wild should also be carefully considered because it may further reduce the effective breeding population.
2. Camera traps provide valuable presence data but are insufficient for fine-scale movement analysis. Satellite telemetry is essential to address this gap, particularly with fewer than 30 individuals remaining. GPS collars would yield critical insights into habitat use, movements, and survival to inform targeted conservation, particularly during the dispersal stage.
3. Road collisions, particularly along the Tehran–Mashhad highway, are now the leading cause of mortality, especially for mobile females and cubs. Urgent mitigation, including fencing, wildlife crossings, and improved road lighting, is required (Mohammadi et al., 2018; 2023).
4. Human removal and illegal capture were documented among the recorded causes of cheetah loss. However, the extent of illegal wildlife trade could not be estimated reliably from our data and requires further investigation as well as anti-trafficking enforcement (Sardari et al., 2026).
5. Cheetah movements extend beyond formal reserves. In the Northern Landscape, unprotected transitional habitats have become vital for breeding and connectivity. Community-based conservation in these areas is essential for long-term persistence (Ahmadi et al., 2017).

The collapse of southern populations illustrates the need to revise protocols to prevent similar losses in the north. While reserves in the Southern Landscape should be maintained for possible dispersal from the Northern Landscape, conservation efforts must prioritize the Northern Landscape, with future reintroductions to the south considered only once population growth is achieved.

## Author contributions

AT, MP and MSF conceived the study, designed the methodology, conducted fieldwork, analysed data, and led writing. MSH, NG, KH, MHK, NS, HA, A. Ranjbaran, A. Radman, AS, and MP contributed to fieldwork, data collection, and validation. MSF supervised the project and reviewed drafts. All authors gave final approval for publication and agreed to be accountable for the work.

## Acknowledgement

We thank the Iranian Department of Environment for collaboration and access to data. We also acknowledge the conservation efforts by the Conservation of Asiatic Cheetah Project (CACP) to protect Asiatic cheetah habitats since 2001. We are grateful to Stichting SPOTS and other funders for their generous support. MSF was supported by Research England’s Expanding Excellence in England (E3) Fund and UK Research and Innovation. We also thank biologists, rangers, citizen scientists, and social media contributors who shared photographic records of Asiatic cheetahs, used in accordance with applicable laws and ethical standards. Finally, we acknowledge the rangers of the surveyed reserves for their invaluable field assistance and data collection.

## Ethical standards

This study was conducted in accordance with guidelines for wildlife research and followed the ethical standards outlined by Oryx for camera-trapping studies. Camera traps were deployed solely for the purpose of monitoring wildlife and were positioned to minimise the likelihood of capturing images of people, avoiding roads, footpaths, settlements, and other areas of regular human activity as far as possible. A small number of images of people were inadvertently obtained which were not used for analysis. All camera-trap surveys were conducted with the permission of the Iranian Department of Environment under permit number 98/420/50475, ص/1400/2074. No animals were handled or disturbed as part of this study, and camera trapping was conducted as a non-invasive monitoring method.

## Data availability

The data supporting this study are available from the corresponding author upon reasonable request, subject to restrictions imposed by the Iranian Department of Environment (DoE).

## Supplementary Materials

**Table S1.** Details of documented losses from the wild cheetah population in Iran during 2012–2024, including deaths and removals from the wild. “Outcome” distinguishes deaths from removals, whereas “Cause or circumstances” describes the reported cause of death or the circumstances of removal. A = adult; S = subadult. In the landscape column, N = Northern Landscape and S = Southern Landscape. Records of deaths and removals from the wild were compiled from reports provided or published by the Iranian Department of Environment.

| Year | Number of individuals | Cause or circumstances | Outcome | Adult/Subadult | Northern/Southern Landscape |
|---|---|---|---|---|---|
| 2012 | 1 | Human-wildlife conflict | Death | A | N |
| 2013 | 2 | Human-wildlife conflict | Death | A | N |
| 2014 | 2 | Road collision | Death | A/S | N |
| 2015 | 1 | Unknown | Death | A | N |
| 2016 | 1 | Human-wildlife conflict | Death | A | N |
|  | 1 | Road collision | Death | A | N |
| 2017 | 1 | Illegal trade | Removal from the wild | S | N |
| 2018 | 2 | Road collision | Death | A/S | N |
|  | 1 | Unknown | Death | A | S |
| 2019 | 0 | _ | _ |  | _ |
| 2020 | 0 | _ | _ |  | _ |
| 2021 | 1 | Unknown | Death | A | N |
| 2022 | 2 | Taken by a shepherd | Removal from the wild | S | N |
|  | 1 | Road collision | Death | S | N |
| 2023 | 2 | Road collision | Death | A/S | N |
|  | 1 | Veterinary malpractice | Death | A | N |
|  | 1 | Natural causes | Death | A | S |
| 2024 | 1 | Malnourishment | Removal from the wild | S | N |

